# Vitamin D and the ability to produce 1,25(OH)_2_D are critical for protection from viral infection of the lungs

**DOI:** 10.1101/2022.06.29.498158

**Authors:** Juhi Arora, Devanshi Patel, McKayla J. Nicol, Cassandra J Field, Katherine H. Restori, Jinpeng Wang, Nicole E. Froelich, Bhuvana Katkere, Josey A. Terwilliger, Veronika Weaver, Erin Luley, Kathleen Kelly, Girish S. Kirimanjeswara, Troy C. Sutton, Margherita T. Cantorna

**Author notes:** Corresponding authors Margherita T. Cantorna, Troy C. Sutton. These authors contributed equally to the manuscript and are listed randomly (Devanshi Patel, McKayla J. Nicol, Cassandra J Field). Funding of the work NIH R01AT005378 to MTC, R01 AI 123521 to GSK, and T32GM108563 to JA, USDA NIFA award PEN04771 to MTC, TCS and GSK.

## Abstract

Vitamin D supplementation has been linked to improved outcomes from respiratory virus infection, and the COVID19 pandemic has renewed interest in understanding the potential role of vitamin D in protecting the lung from viral infections. Therefore, we evaluated the role of Vitamin D using animal models of pandemic H1N1 influenza and SARS-CoV-2 infection. In mice, dietary induced vitamin D deficiency resulted in lung inflammation that was present prior to infection. Vitamin D sufficient (D+) and deficient (D-) wildtype (WT) and D+ and D-Cyp27B1 (Cyp) knockout (KO, cannot produce 1,25(OH)_2_D) mice were infected with pandemic H1N1. D- WT, D+ Cyp KO, and D- Cyp KO mice all exhibited significantly reduced survival compared to D+ WT mice. Importantly, survival was not the result of reduced viral replication as influenza M gene expression in the lungs was similar for all animals. Based on these findings, additional experiments were performed using the mouse and hamster models of severe acute respiratory syndrome coronavirus-2 (SARS-CoV-2) infection. In these studies, high dose vitamin D supplementation reduced lung inflammation in mice but not hamsters. A trend to faster weight recovery was observed in 1,25(OH)_2_D treated mice that survived SARS-CoV-2 infection. There was no effect of vitamin D on SARS-CoV-2 N gene expression in the lung of either mice or hamsters. Therefore, vitamin D deficiency enhanced disease severity, while vitamin D sufficient/supplementation reduced inflammation following infections with H1N1 influenza and SARS-CoV-2.

## Introduction

Low vitamin D status has been associated with poorer outcomes following acute respiratory diseases including influenza [1]. Vitamin D supplements have been touted as being useful in high doses for reducing the severity of seasonal influenza [2-4]. The recent emergence of severe acute respiratory syndrome (SARS)-coronavirus (CoV)-2 and the ongoing pandemic has led to a renewed interest in high dose vitamin D supplements to prevent and treat severe SARS-CoV-2 disease (i.e. COVID-19) [4]. Infection with SARS-CoV-2 results in local and systemic inflammation that when controlled may enhance survival and clinical outcomes of COVID-19 [5]. Severe respiratory illness can also be caused by influenza viruses or co-infection with influenza and coronaviruses. An association between low vitamin D status and severe COVID-19 has been postulated and accordingly, vitamin D supplementation has been proposed to be beneficial to treat COVID-19 [6]. A recent systemic review concluded that low circulating levels of vitamin D (serum 25(OH)D, 25D) were associated with more severe symptoms, and higher mortality in patients with COVID-19 [7]. Interventions that control the lung inflammatory response to viruses have the potential to benefit the global population.

Vitamin D, and the active form of vitamin D (1,25(OH)_2_D, 1,25D) has been implicated to play a role in the anti-viral response; however, this effect may be specific to different viruses. For example, 1,25D treatment of T cells from human immunodeficiency virus (HIV) infected individuals *in vitro* resulted in a decrease in viral RNA transcription by direct reduction of NF-κB, which reactivates proviral HIV [8]. Conversely, 1,25D treatment of respiratory syncytial virus (RSV) infected human epithelial cells *in vitro* did not affect viral replication [9]. Production of antimicrobial peptides such as cathelicidin, β-defensin etc., and production of cytokines such as TNFα, IL-5, IL-1β, IL-6, IL-10 and type I interferons at the mucosal surface are important parts of the innate immune response against viruses [10,11]. Several studies have shown that 1,25D and other vitamin D analogs induced cathelicidin production in response to virus infection [12]. Cathelicidin LL-37 has been shown to bind and kill viruses including influenza viruses *in vitro* [13-17]. Therefore, it is possible that vitamin D through the induction of cathelicidin could directly target SARS-CoV-2 and influenza. 1,25D also limits inflammatory responses by decreasing IFNγ, IL-6 and TNFα [18,19]. The effects of 1,25D to reduce inflammation could be detrimental for the ability of the host to clear some viruses.

Lung epithelial cells are vitamin D targets since they express the vitamin D receptor (VDR) and are regulated by 1,25D treatments. In mice, 1,25D treatments reduced inflammation following lipopolysaccharide induced lung injury and regulated angiotensin converting enzyme (ACE) expression in the rat lung epithelium [20]. In mice, infection with an influenza H9N2 virus induced mRNA for the VDR in the lung and 1,25D treated animals had reduced lung inflammation [21]. Treating mice with the 1,25D precursor, 25hydroxyvitamin D (25D), also protected mice from subsequent H1N1 influenza infection [22]. 1,25D treatment of human bronchial epithelial cells suppressed IL-6 and protected the cells from oxidative damage [23]. VDR knockout (KO) mice had reduced expression of tight junction proteins such as zonula occludens-1, occludin, and claudins (2,4, and 12), suggesting an important role for the VDR in maintaining the integrity of the lung [24]. Vitamin D has direct effects on the lung epithelium and 1,25D suppresses inflammation in the lung.

Extensive association studies in humans have led to the proposal that high dose vitamin D supplements could be beneficial for protecting from severe influenza and SARS-CoV-2 infections. Since cause and effect are extremely difficult to determine in human studies: we sought to evaluate the effects of vitamin D on the lung anti-viral response in animal models. The data from mice and hamsters suggests that vitamin D supplementation reduces inflammation in the lung following pandemic H1N1 and SARS-CoV-2 infection. D-mice had lung inflammation even without infection. We had shown previously that feeding D-diets to mice that cannot produce 1,25D (Cyp KO) resulted in severe vitamin D deficiency [25]. Influenza disease was greatest in D- Cyp KO and the least amount of disease was in D+ WT mice. The survival of D+ Cyp KO mice was significantly less than D+ WT mice following an influenza infection. Vitamin D supplementation reduced lung inflammation and *Ifnβ* expression in the lung of mice following SARS-CoV-2 infection. Vitamin D treatments had no effect on the expression of viral RNA for either SARS-CoV-2 or H1N1 influenza in the lungs of hamsters or mice. Instead, the data support an important role for vitamin D and 1,25D in controlling the host inflammatory response to viruses in the lungs.

## Materials and Methods

### Animal models

C57BL/6 WT, K18hACE2 (Jackson Laboratories, Bar Harbor, ME) and Cyp KO (gift from Dr. Hector DeLuca, University of Wisconsin, Madison, WI) mice were bred (mice) and housed (mice and hamsters) according to approved IACUC protocols at the Pennsylvania State University (University Park, PA). For experiments, age and sex-matched mice were fed: Chow diets (D+) (Lab diets # 5053, Arden Hills, MN) or purified diets with (D+) and without (D-) vitamin D (Envigo, T.D. 89123, Madison, WI). For some experiments D+ mice were fed corn oil alone or corn oil with 1,25D. For some experiments D-mice were fed corn oil alone or corn oil with one of two doses of vitamin D3 (Sigma-Aldrich, C9756, St. Louis, MO). Age and sex-matched Golden Syrian Hamsters were purchased from Envigo (Indianapolis, IN) and maintained on chow (D+) or D-diet (Envigo, T.D.120008) and orally fed corn oil or corn oil with vitamin D3. Serum was collected to monitor vitamin D status of mice and hamsters.

### Serum 25hydroxy vitamin D (25D) measurements

Serum 25D levels were measured using an ELISA kit and standards as per the manufacturer’s instructions (25-OH D, Eagle Biosciences, Amherst, NH). The limits of detection were 1.6 ng/ml 25D.

### SARS-CoV-2 infection

SARS-CoV-2 USA-WA-1/2020 (Centers for Disease Control and Prevention, BEI Resources, NIAID, NIH: NR-52281) was used for infecting both mice and hamsters (The World Reference Center for Emerging Viruses and Arboviruses, University of Texas Medical Branch, Galveston TX, at passage 4). K18hACE2 mice were anesthetized using isofluorane and infected with 100-1000 TCID_50_ units of SARS-CoV-2 in 50 µL phosphate buffered saline. Hamsters were sedated with ketamine (150 mg/kg), atropine (0.015 mg/kg) and xylazine (7.5 mg/kg) via intraperitoneal injection and intranasally inoculated with 10000 TCID_50_ units of SARS-CoV-2 in 100 µL Dulbecco’s Modified Eagle Media. Hamsters were given atipamezole (1 mg/kg) subcutaneously to reverse the sedation. Equal numbers of males and females were used for all experiments and body weights and symptoms were monitored daily until end-point criteria or day 14 post-infection.

One series of experiments in K18hACE2 mice used standard D+ rodent chow diet (Lab diets # 5053, Arden Hills, MN) diets with oral dosing of corn oil or 10ng/d of 1,25D diluted in 10µl of corn oil beginning the day before infection and continuing until sacrifice. Additional experiments used mice or hamsters fed D-diets with or without oral vitamin D3. D-K18hACE2 mice were dosed orally with corn oil (D-), 0.125 µg vitamin D3/d, (D+) or 2.5 µg vitamin D3/d (D++) beginning 8 weeks prior to infection and continuing throughout the experiment. Hamsters were placed on D-diets with corn oil or 8 µg vitamin D3 (D+) in corn oil/d starting 11 days before infection and continuing throughout the experiment.

### H1N1 infection

D+ and D- WT and D+ and D- Cyp KO littermates were fed identical diets with and without vitamin D. Mice were anesthetized with isofluorane to intranasally inoculate them with 10 – 30 TCID_50_ mouse-adapted A/H1N1/California/04/2009 influenza in 50 µL Gibco’s reduced-serum minimum essential media (ThermoFisher, 31985-070, Waltham, MA) [26]. Body weight and clinical signs were monitored daily until they either reached end point criteria or at d14 post-infection.

### Biocontainment and Animal Care and Use

All studies with SARS-CoV-2 were conducted in a biosafety level 3 enhanced (BSL3+) laboratory. This facility is approved for BSL3+ respiratory pathogen studies by the US Department of Agriculture and Centers for Disease Control. Studies with pandemic H1N1 influenza were conducted under biosafety level 2 enhanced conditions. All animal studies were conducted in compliance with the Animal Care and Use Committee under protocol numbers: 202001693, 202001516, 202001440, and 202001638.

### RNA isolation and quantitative PCR

Tissues were homogenized in TRIzol reagent (Sigma-Aldrich, St. Louis, MO). Total RNA was extracted from the tissues using chloroform-isopropanol precipitation and quantified using NanoDrop (ThermoFisher, Waltham, MA). 1-2 µg RNA was reverse transcribed using AMV Reverse Transcriptase (Promega, Madison, WI). Primers were purchased from Integrated DNA Technologies (IDT, Coralville, IA) and are listed in Supplementary (S)Table. 1. Gene expression was quantitated using SYBR green mix (Azura Genomics, Raynham, MA) and StepOne Plus system (Applied Biosystems, Carlsbad, CA). Primers for SARS-CoV2 N gene were a commercial kit from IDT (Cat# 10006713). Gene expression was calculated using the deltadelta Ct method using GAPDH and uninfected control tissues. Gene expression was normalized to day 0 uninfected controls.

### Histology

Lung tissues were collected from hamsters and mice and fixed in 10% formalin. These tissues were embedded in paraffin, sectioned, and H&E stained by the PSU Animal Diagnostic Lab Histology lab. Scoring of lung pathology associated with SARS-CoV-2 infection was done by veterinary pathologists certified by the American College of Veterinary Pathologists that were blinded to vitamin D3 treatment status. The criteria for the histopathologic evaluation of the tissue sections was done as described for H1N1 infected mouse lungs [27-29], SARS-CoV-2 infected mouse lung sections [30,31], and hamster lung sections [32-34]. The individual parameters evaluated the extent of pneumonia, damage to alveoli and lymphocytic infiltration by various specific parameters described in STable 2-4.

### Statistical analysis

Results are represented as mean ± SEM. Statistical analysis was performed using Prism ver. 10 (GraphPad, San Diego, CA) using Multiple t-tests, One-way ANOVA, Two-way ANOVA, Mixed-effects analysis with Bonferroni multiple comparisons tests, Kruskal-Wallis test with Dunn’s multiple comparison, Unpaired t-test, and Log-rank survival test as applicable. Data was checked for normal distribution and outliers. P < 0.05 was the cutoff to determine significance.

## Results

### The effect of vitamin D on H1N1 infection

To evaluate the role of vitamin D deficiency during respiratory virus infection, WT and Cyp KO mice were fed a D+ or D-diet. Regardless of the genotype, serum 25D levels were significantly different between mice fed D+ or D-diet (Fig. 1A). As expected D+ Cyp KO mice accumulated 25D and had higher levels of 25D than D+ WT mice [25]. Following infection with pandemic H1N1 influenza, respiratory distress was evident in all mice by 7 days post-infection (Fig. 1B). D+WT mice showed the least amount of respiratory distress and after day 10 D+ WT and D+ Cyp KO mice no longer exhibited symptoms of respiratory distress (Fig. 1B). D- WT and D- Cyp KO mice had greater symptoms of respiratory distress that were not completely resolved by d14 post-infection in the D- Cyp KO mice (Fig. 1B). Only 1 of 18 D+ WT mice died following H1N1 infection (Fig. 1C). D- WT mice had lower survival compared to D+ WT mice (Fig. 1C). Conversely, both Cyp KO groups showed decreased survival; 62% from D+ Cyp KO and 57% from D- Cyp KO mice, respectively. The expression of the influenza M gene at d4 post-infection was not different in the 4 groups of influenza infected mice (data not shown). Lung sections from D- WT and D- Cyp KO mice (d0) but not D+ WT mice showed signs of inflammation even before infection (Fig. 1D). The amount of alveolar hemorrhage was significantly more at d4 post-infection in the D- Cyp KO compared to D+ WT or D- WT (Fig. 1D). At d14 post-infection the lung sections of D+ Cyp KO, D- WT and D- Cyp KO mice appeared more severe than D+ WT (Fig. 1E). Therefore, Vitamin D deficiency and the inability to produce 1,25D increased the susceptibility of mice to H1N1 influenza infection.

**Figure 1.**
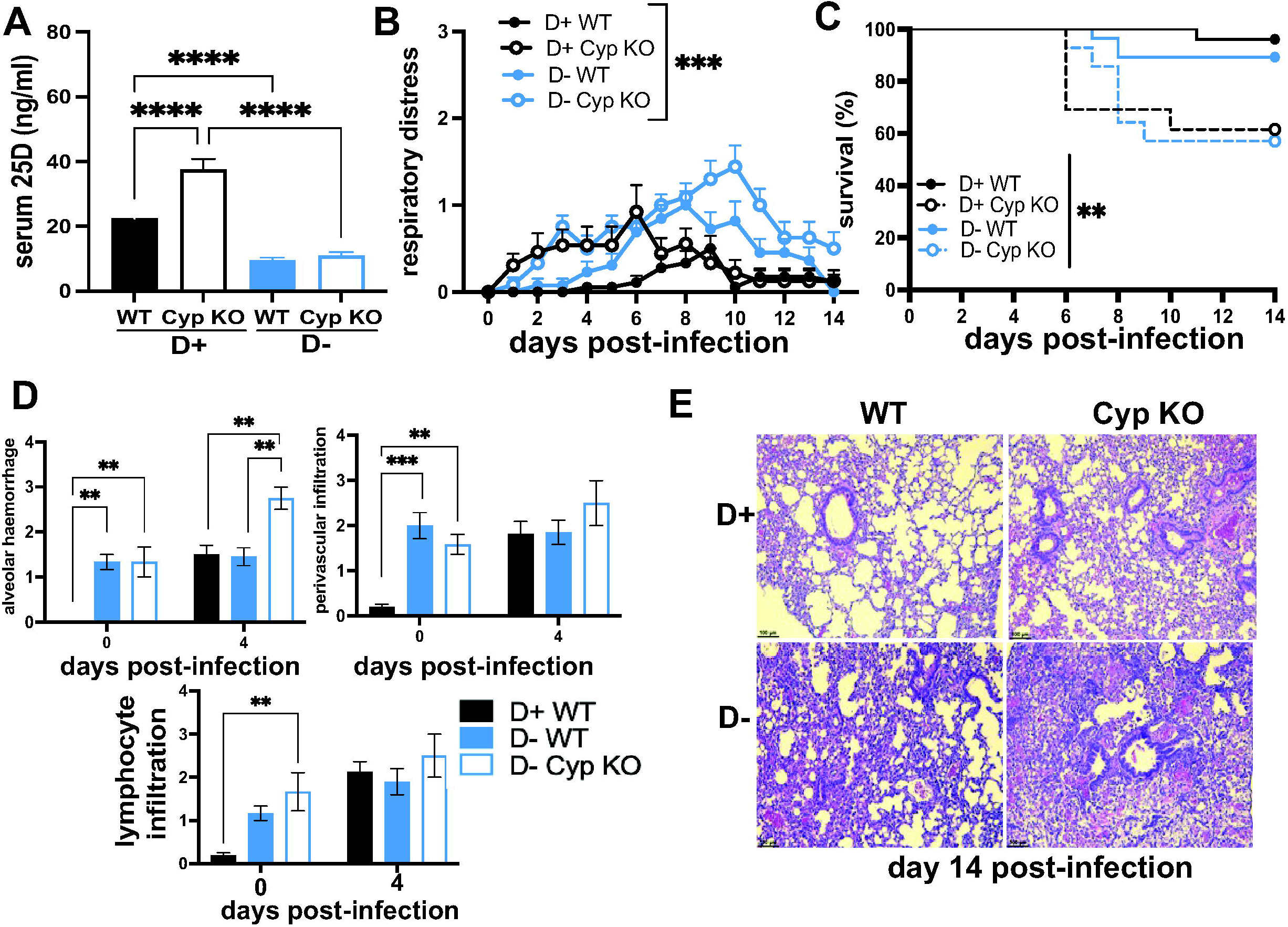
Vitamin D deficiency and Cyp27B1 KO increases susceptibility to H1N1 infection. D+ and D- WT and D+ and D- Cyp KO littermates were infected with H1N1 influenza (n=13-18 per group). **A)** Serum 25D, **B)** respiratory distress symptoms and **C)** survival of D+ WT (n=18), D+ Cyp KO (n=13), D- WT (n=13) and D- Cyp KO (n=12) mice at d14 post-infection. **D)** lung alveolar hemorrhage, perivascular infiltration and lymphocyte infiltration in D+ WT (n=4) and D- WT (n=4-5) and D- Cyp KO (n=2-3) mice at d0 and d4 post-infection. **E**) Representative histology of the lung of D+ WT (score = 3), D+ Cyp KO (score = 4), D- WT (score = 5.5) and D- Cyp KO (score = 6) at d14 post-infection. Values are the mean ± SEM. Statistical significance was assessed using one-way ANOVA with Bonferroni multiple comparison test for **(A)**, two-way ANOVA with Bonferroni multiple comparison test for **(B & D)** and Log rank (Mankel-Cox) survival analysis for **(C)**. **P < 0.01, ***P < 0.001 and ****P < 0.0001.

### Mouse and hamster models of SARS-CoV-2 infection

SARS-CoV-2 do not effectively infect WT mice. The transgenic expression of human (h)ACE-2 has been shown to allow SARS-CoV-2 infection in mice [35]. hACE-2 mice were infected with 100, 200, and 315 TCID_50_ SARS-CoV-2 and the mice were evaluated for pre-determined euthanasia endpoints for up to 14d post-infection. The dose of SARS-CoV-2 that resulted in the sacrifice of 50% of the mice was 200 TCID_50_ (Fig. 2A). The mice infected with 100 TCID_50_ failed to reach the euthanasia endpoints, were sacrificed at d14 post-infection and had only minimal weight loss (Fig. 2A). The viral gene copy number for the nucleocapsid (N) protein was measured in the lungs of mice at 100 and 200 TCID_50_. High copy numbers of the N gene were detected as early as d2 post infection and remained high until d6 post infection (Fig. 2B). The N gene copy number significantly decreased by d14 post-infection (Fig. 2B). Interestingly, the mice that were infected with 100 TCID_50_ had high amounts of the N gene in the lung even though they showed only mild symptoms of infection and there were no differences between the N gene expression in the lung between the 100 and 200 TCID_50_ inoculum (Fig. 2A and Fig. 2B). Lungs from SARS-CoV2 infected mice were used to measure mRNA for the 1alpha hydroxylase that produces 1,25D (*Cyp27B1*), vitamin D receptor (*Vdr*), and the 24hydroxlase that degrades vitamin D (*Cyp24A1*). There was no change in *Vdr* expression in the lung at d2 or d4 post-infection (Fig. 2C). Conversely, *Cyp27B1* and *Cyp24A1* expression was higher at d2 and d4 post-infection than in the uninfected lung (Fig. 2C). *Cyp27B1* expression was significantly lower in the d4 than the d2 post-infection lung (Fig. 2C). Infection of the hACE2 mice with SARS-CoV-2 induced the expression of two genes that regulate vitamin D metabolism in the lungs.

**Figure 2.**
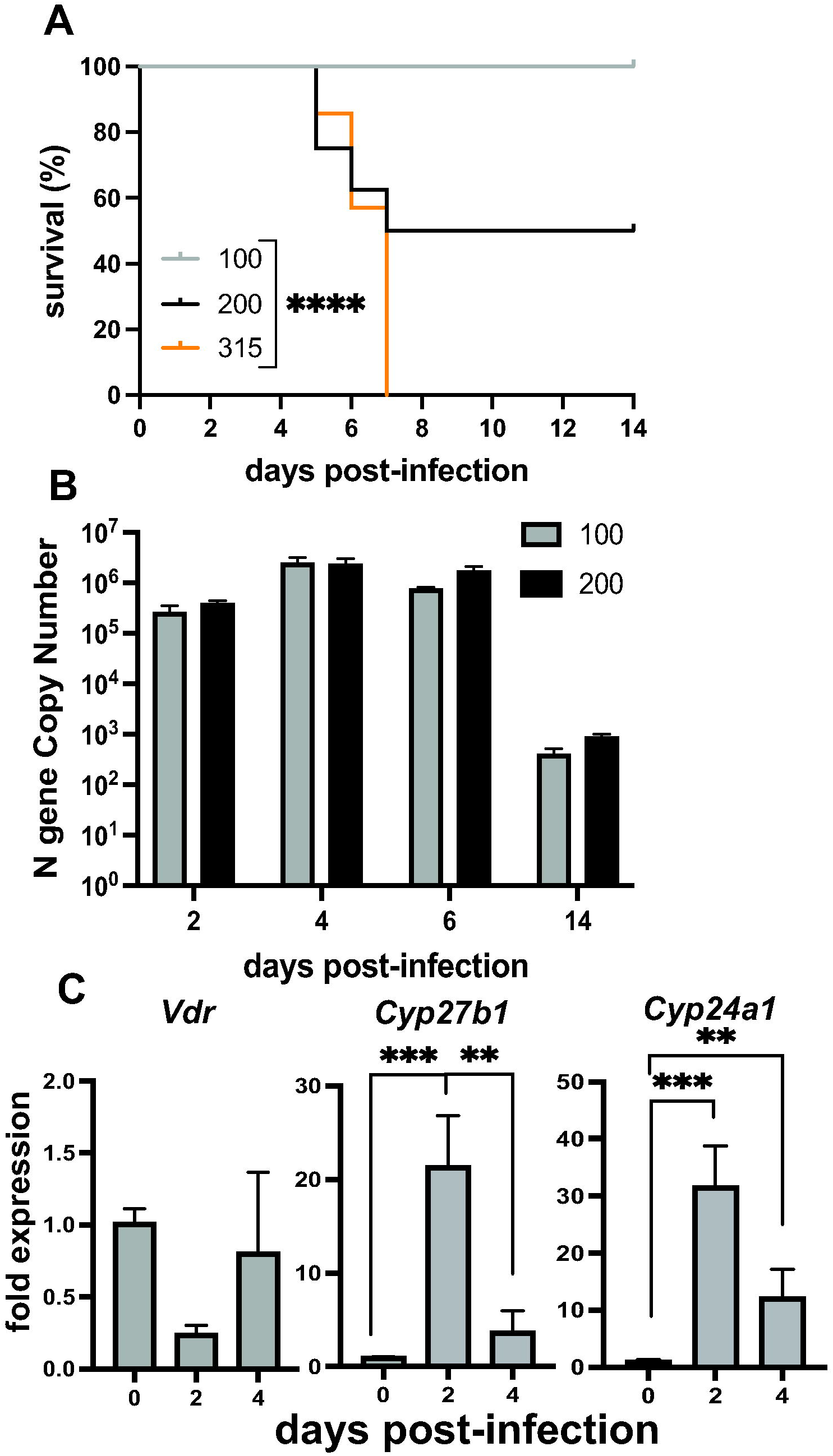
SARS-CoV-2 infection. K18-hACE2 mice were intranasally inoculated with 100 (n=8), 200 (n=8) and 315 (n=7) TCID_50_ of SARS-CoV-2 virus. Mice were sacrificed when they met pre-determined endpoints or at d14 post-infection. **A)** Survival, **B)** SARS-CoV-2 N gene expression in the lung, **C)** *Vdr, Cyp24A1* and *Cyp27B1* mRNA gene expression (n = 4/group/timepoint) at d0, d2 and d4 post-infection in the lung. Samples were normalized to uninfected control tissue. Values are the mean ± SEM. Statistical significance was assessed using Log-rank test for trend (**A**), two-way ANOVA model for main effects only for (**B**) and one-way ANOVA with Bonferroni multiple comparison test on log-transformed expression values for (**C**). *P < 0.05, ***P < 0.001 and ****P< 0.0001.

Hamsters infected with SARS-CoV-2 are reflective of human SARS-CoV2 infection, in that the virus infects the lower respiratory tract, causing similar respiratory sequelae [36]. We have previously shown that hamsters infected with 10^5^ TCID_50_ SARS-CoV-2 lost weight shortly after infection and none of the hamsters died following infection [33]. The weight loss peaked by d5 post-infection, and the hamsters recovered completely by d10 post-infection [33].

### The effect of vitamin D on SARS-CoV-2 infection

To establish differing Vitamin D status, ACE2 mice were given a vitamin D deficient chow and were orally dosed with vehicle (D-), 0.125µg/d (D+) or 2.5µg/d (D++) for 8wks prior to infection. Serum 25D levels were higher in D+ mice and significantly higher in D++ mice before and 14d post-infection (Fig. 3A). The D+ dose was inadequate to significantly raise serum 25D levels over the D-values (Fig. 3A). The survival of the SARS-CoV2 infected mice was not affected by the D+ or the D++ treatments (Fig. 3B). At d6 post-infection the amount of N gene expression was the same in D- and D++ mice (Fig. 3C). The expression of the *Vdr, Cyp27B1, Cyp24A1* were not different in D- and D++ mice at d6 post-infection (Fig. 4A). *Ifnβ* and *Ifnγ* were induced by SARS-CoV2 infection, while *Ifnα* was not (uninfected control set at 1, Fig. 4B). Expression of *Ifnβ* was significantly lower in the D++ lung as compared to the D-lung at d6 post-infection (Fig. 4B). Mice surviving until d14 post-infection showed no difference in lung histopathology scores between D- and D+ mice (Fig. 3D&E). The D++ lung histopathology scores showed significantly reduced type II hyperplasia, significantly reduced alveolar remodeling and lower (not significant) total histopathology scores (Fig. 3D&E). The final series of experiments tested whether the active form of vitamin D (1,25D) could prevent the lethality of SARS-CoV-2 infection. There was no effect of 1,25D on survival from a lethal dose of SARS-CoV-2 (1000 TCID_50_) (Fig. 3F). There was a trend for faster weight recovery in the surviving 1,25D treated mice that did not reach significance (Fig. 3F). There was an effect of D++ treatment to decrease lung histopathology and a trend towards faster recovery in 1,25D treated mice infected with SARS-CoV-2.

**Figure 3.**
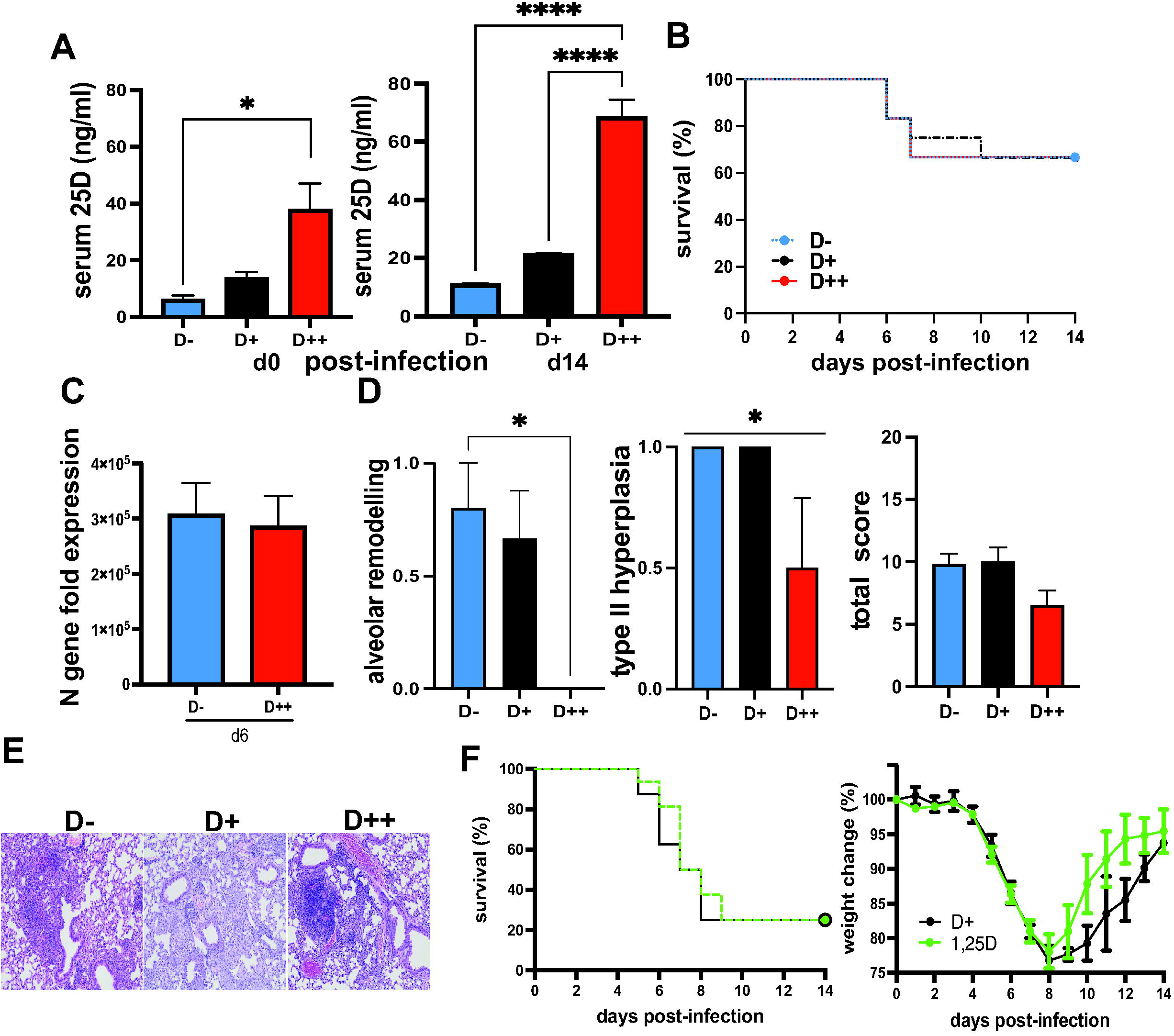
High dose vitamin D reduces lung inflammation following SARS-CoV-2 infection. hACE2 mice were fed vitamin D deficient (D-, (n=18), vitamin D sufficient (D+, n=12) or vitamin D supplemented (D++, n=18) diet and infected with 200 TCID_50_ SARS-CoV-2. **A)** Serum 25D was measured before and d14 post-infection, and **B**) the survival of D-, D+, and D++ mice. **C**) SARS-CoV-2 N gene expression in the lungs (n=6 mice/group) at d6 post-infection. Gene expression relative to uninfected D+ controls. **D)** Alveolar remodeling, type II pneumocyte hyperplasia and total histology score in D-, D+ and D++ mice (n=4-6 mice /group) at d14 post-infection. **E)** Representative histology images for D-(score = 9), D+ (score = 11) and D++ (score = 5). D+ hACE2 (n=8) mice at d14 post-infection. 1,25(OH)_2_D (1,25D) treated D+ hACE 2 (n=16) mice were infected with 1000 TCID_50_ SARS-CoV-2. **F)** Survival and body weight change over the course of infection. Values are the mean ± SEM. Statistical significance was assessed using one-way ANOVA with Bonferroni multiple comparison test for (**A & D)**, Log rank (Mankel-Cox) test for each of the groups for (**B & F)**, Unpaired t-test on log-transformed expression values for (**C)** and two-way ANOVA with Bonferroni multiple comparison test for (**F)**. *P < 0.05 and ****P < 0.0001.

**Figure 4.**
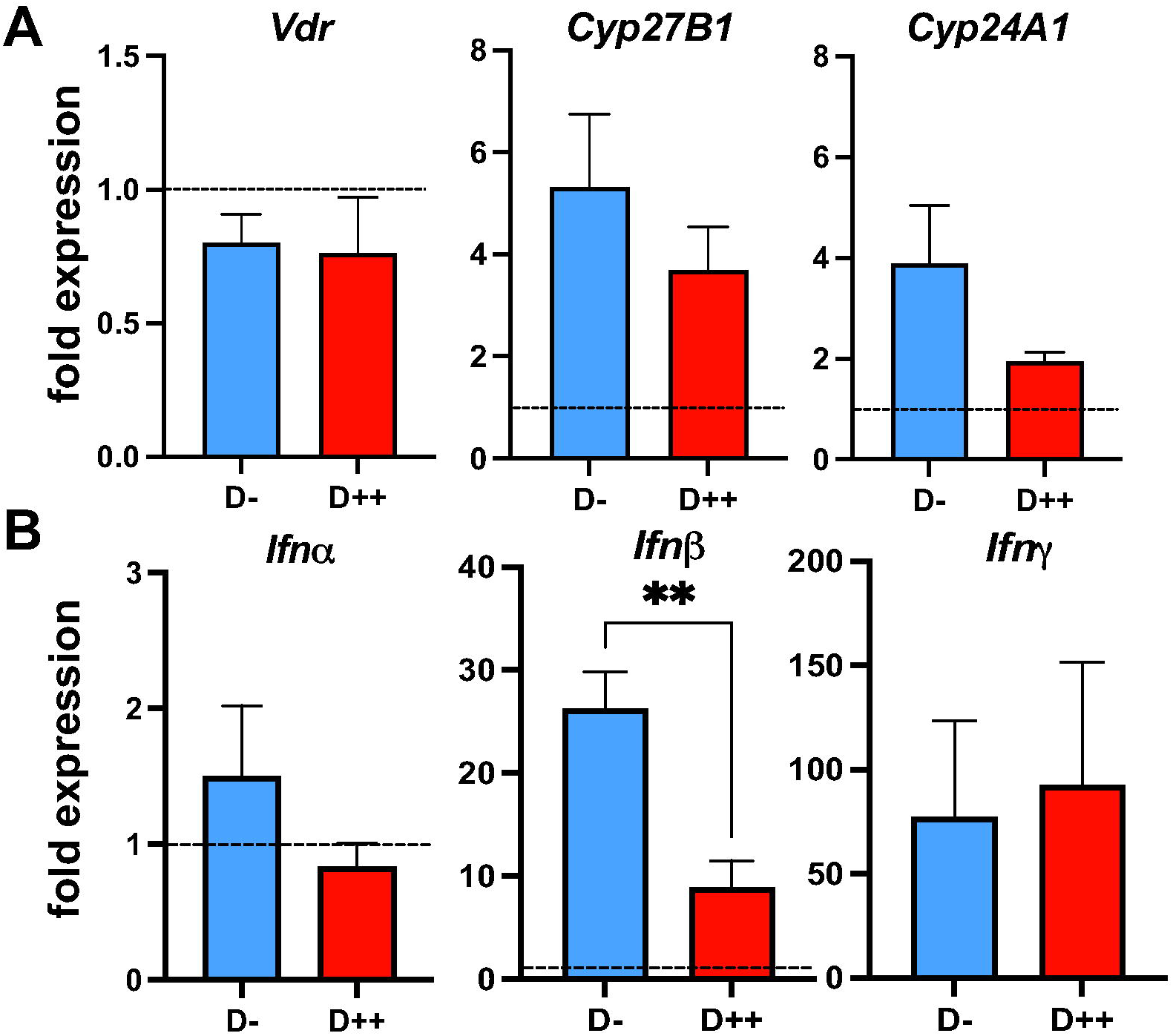
D++ mice have lower *Ifnβ* at d6 post-SARS-CoV-2 infection in the lung. Lung mRNA from d6 SARS-CoV-2 infected D-(n=5) and D++ (n=5) mice. **A)** *Vdr, Cyp27B1, Cyp24A1* and **B)** *Ifnα, Ifnβ* and *Ifnγ* relative to *Gapdh* and uninfected controls set at 1 (dashed line). Values are the mean ± SEM. Statistical significance was assessed using unpaired t-test on log-transformed values. **P<0.01.

Serum 25D levels in hamsters fed on chow diet were not significantly different than serum 25D levels in hamsters fed D-diets for 4 weeks (Fig. 5A). Therefore, to control for the diet, hamsters were fed D-diets and then fed orally with vehicle (D-) or with 8µg/d of vitamin D3 (D+) beginning 14 days before the SARS-CoV-2 infection and continuing throughout the experiment. Confirming the effectiveness of the dietary intervention, the serum 25D levels were significantly higher in D+ hamsters as compared to D-hamsters before infection on d0 and this effect was maintained until d14 post-infection (Fig. 5A). SARS-CoV-2 N gene was detected on d3 but not at d6 post-infection in the lungs (Fig. 5B) and there was no difference in N gene expression between the D+ versus D-lungs (Fig. 5B). Surprisingly, SARS-CoV-2 N gene expression was also detected in the colon tissues of hamsters at both d3 and d6 post-infection compared to uninfected control tissues (Fig. 5B). Expression of the N gene in the colon was 1000-fold less than in the infected lung (Fig. 5). N gene expression was significantly higher in the D-colon compared to baseline values from uninfected tissue controls but not different from uninfected control in the D+ colon at d3 post-infection (Fig. 5B). The histopathology of the lungs showed significantly more damage at d6 than d3 post-infection (Fig. 5C, 5D). There was no effect of vitamin D on the weight loss or histopathology scores following SARS-CoV-2 infection of hamsters (Fig. 5D & 5E).

**Figure 5.**
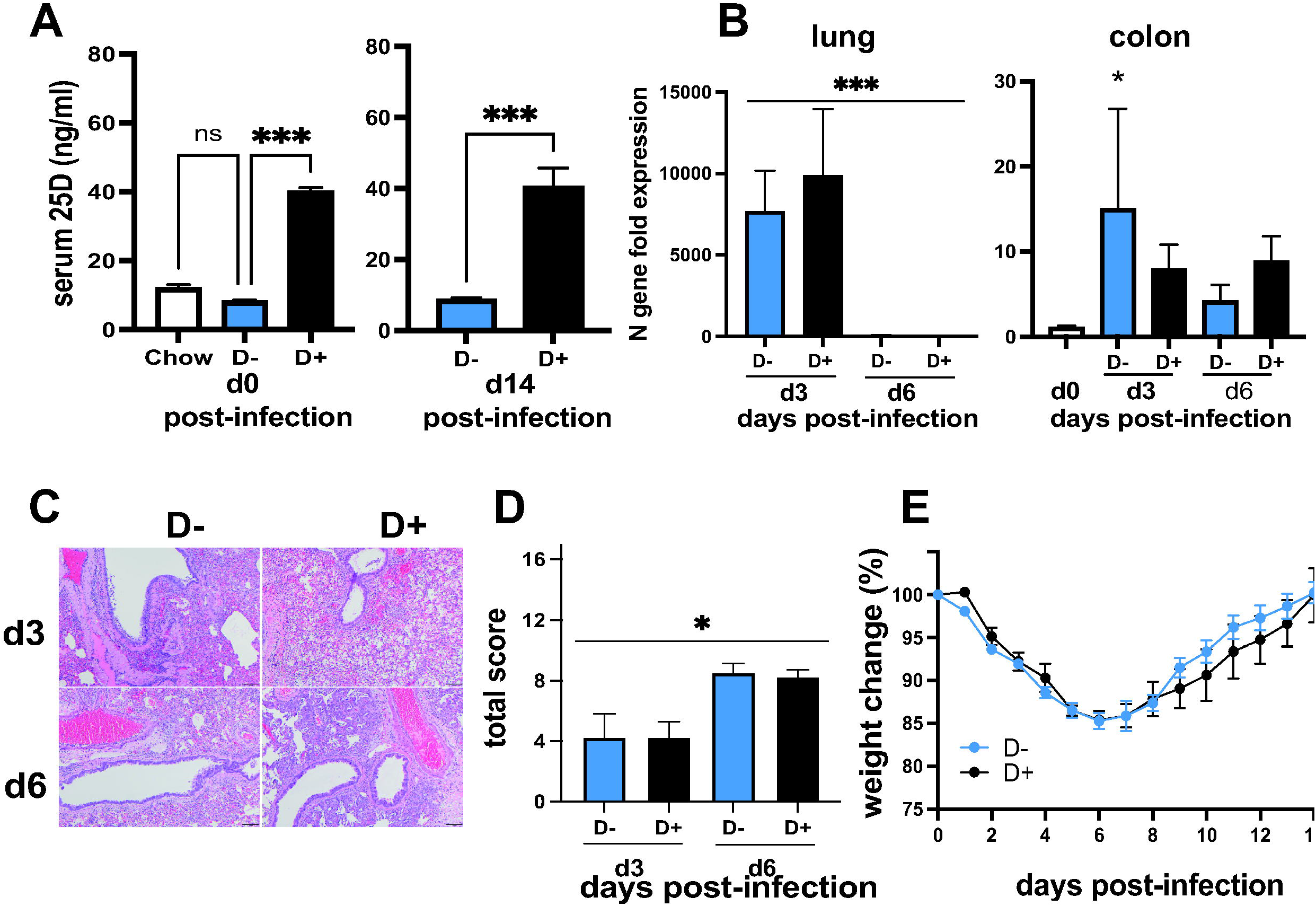
SARS-CoV-2 infection of hamsters. **A**) Serum levels of 25D in chow-fed, D+ and D-fed hamsters before (n=2-10 hamsters/group) or 14d after (n=5-6 hamsters/group) SARS-CoV2 infection. **B**) SARS-CoV-2 N gene expression in the lung and colon relative to uninfected control (n=4 hamsters per group and timepoint). **C**) Representative histopathology sections of the lungs at d3 (D-score = 6, D+ score = 4) and d6 (D-score = 10, D+ score = 9) post-infection and **D)** total histopathology scores (n=4 hamsters/group). **E)** Change in body weight following infection with SARS-CoV2 (n= 5-6 hamsters/group). Values are the mean ± SEM. Statistical significance was assessed using Kruskal-Wallis test with Dunn’s multiple comparison (d0) & Unpaired t-test (d14) for **(A)**, one-way ANOVA with Bonferroni multiple comparison test on log-transformed expression values for **(B)**, one-way ANOVA for **(D)** and two-way ANOVA with Bonferroni multiple comparison test for **(E)**. *P < 0.05 and ***P < 0.001.

## Discussion

Vitamin D deficiency resulted in lung inflammation in the absence of infection. Infected D-mice had more severe lung inflammation and respiratory symptoms than D+ or D++ mice when infected with either H1N1 influenza or SARS-CoV-2. The data point to shared effects of vitamin D to control inflammation in the lung following influenza or coronavirus infection. D-mice had significantly more inflammation than D+ mice following H1N1 influenza infection (Fig. 1). High dose vitamin D treatment (D++) resulted in some protection of mice from SARS-CoV-2 (Fig. 3). In addition, 1,25D treatment showed a trend toward faster recovery of surviving mice from SARS-CoV-2 (Fig. 3). Others have shown that 1,25D treated mice had reduced lung inflammation [20,21], and treating D+ mice with 25D had a small protective effect on weight loss and lethality following H1N1 infection [22]. A recent clinical trial that used 25D in humans showed reduced mortality in hospitalized patients with COVID-19 [37,38]. However, it is unclear whether 25D treatment would be effective in mouse or hamster models of SARS-CoV-2. This is the first study that investigated the effects of vitamin D in animal models of SARS-CoV-2. The data suggest that vitamin D and 1,25D may be effective to protect the lung from SARS-CoV-2. SARS-CoV-2 infection induced *Cyp27B1* and *Cyp24A1* in the lung of mice suggesting a role for vitamin D metabolites in the lung response to SARS-CoV-2 infection (Fig. 2). There are likely shared and unique mechanisms by which vitamin D regulates the host response to influenza versus SARS-CoV-2. A better understanding of the mechanisms by which vitamin D regulates the anti-viral response in the lung to both influenza and coronaviruses is needed to inform clinical studies. Cyp27B1 KO mice cannot produce 1,25D, which induces Cyp24A1 and degrades 25D and 1,25D. Feeding Cyp27B1 KO mice D+ diets results in the accumulation of 25D (Fig. 1A, [25,39]). 25D is a low affinity ligand for the VDR and it has been shown that high amounts of 25D can replace the need for 1,25D for the regulation of calcium homeostasis and osteomalacia [40]. Previous experiments showed that D+ Cyp KO and D+ WT mice cleared a bacterial infection in the gut with similar kinetics [25]. Conversely, D+ Cyp KO mice had higher lethality and more severe inflammation than D+ WT mice when infected with H1N1 influenza (Fig. 1). The effects of Cyp27B1 expression on host-resistance to a bacteria could be different than the effects on host-resistance to a virus. It would be interesting to determine the effect of the Cyp27B1 deletion on host resistance to other respiratory viruses including SARS-CoV-2. Conversely the differential effect of Cyp27B1 could be due to the location of the infection in the gut versus the lung. Regardless, it seems that the ability of the host to produce Cyp27B1 is important for the mice to survive an H1N1 lung infection.

The dietary interventions to generate D-, D+ and D++ K18hACE2 mice resulted in D++, but not D+, mice having higher serum 25D than D-mice (Fig. 4). Interestingly, the hamster studies suggest that the commercially available chow may not be adequate to raise serum 25D levels (Fig. 5A). The data suggest that the adequacy of vitamin D should be considered in evaluating studies that infect chow fed hamsters with SARS-CoV-2. At d6 post-SARS-CoV-2 infection, the D++ K18-hACE2 mice had less inflammation and lower IFNβ in the lung than the D-mice (Fig. 3). The results are consistent with the anti-inflammatory effects of vitamin D. Suppression of type-1 inflammatory cytokines by vitamin D underlie the effects of vitamin D and 1,25D to suppress immune mediated diseases [41-43]. The benefits of 25D from influenza infection was associated with a reduction in IFN-γ in the lung [22]. Recently, Chauss et.al showed that 1,25D promotes anti-inflammatory responses by switching off IFNγ production from Th1 cells and upregulating IL-10 [44]. IFN-γ and IFNβ production is essential for effective viral clearance, viruses have mechanisms to evade the IFN responses, and severe COVID-19 is associated with dysregulation of IFN responses [45-47]. Down-regulation of IFNβ by vitamin D is associated with protection from inflammation in the lung following a virus infection with either influenza or SARS-CoV-2. Importantly, there was no effect of vitamin D on SARS-CoV-2 N gene or H1N1 M gene expression in the lungs of mice or hamsters. This indicates that vitamin D did not reduce or inhibit viral replication in the lung. We found SARS-CoV-2 N gene expression in the hamster colons. Interestingly, D-colons had relatively more SARS-CoV-2 N gene expression than D+ colons. The implications of having SARS-CoV-2 in the colon but not the lung would need to be determined and it would be important to quantitate live virus in the tissues. Unfortunately, we did not save colons from our SARS-CoV-2 mouse studies. Vitamin D has been shown to be a strong inducer of cathelicidin LL-37 in human cells [48]. There has been some evidence that LL-37 can directly kill some viruses including influenza viruses [13-17]. Treating mice with a high dose (500µg/d) of human LL-37 peptide protected from lethal influenza infection and significantly reduced viral titers at d3 post-H1N1 infection in the lung [49]. LL-37 inhibited binding of SARS-CoV-2 spike protein containing pseudo viruses both *in vivo* and *in vitro* blocking entry via ACE2 [50]. There were no effects of vitamin D *in vivo* on the expression of viral genes for SARS-CoV-2 or H1N1 influenza in the lung. The cathelicidin peptides found in mice are not the same as the LL-37 in humans and the mouse cathelicidin is not regulated by vitamin D [51]. The lack of a vitamin D effect on SARS-CoV2 was shown in mice and hamster lung. It is unclear whether the hamster cathelicidin gene has vitamin D response elements. Furthermore, no studies have been done to test the effect of vitamin D on SARS-CoV-2 *in vitro*. Therefore, an effect of vitamin D through the induction of anti-bacterial peptides, like LL-37, that reduces viral titers cannot be ruled out.

## Conclusions

Together the data support an important role for vitamin D and Cyp27B1 in the regulation of the host response to H1N1 and SARS-CoV-2 viruses. The role of vitamin D includes the restraining of the IFN response shortly after infection. Vitamin D deficient hosts had pre-existing inflammation in the lungs that contributed to susceptibility to viral infection. Future experiments should continue to determine the mechanisms by which vitamin D regulates the anti-viral response in the lungs and whether there are differences in the effect of vitamin D on host resistance to H1N1 influenza and SARS-CoV-2.

## Supplementary material

STable 1: Primer sequences for qPCR, STable 2: Histological evaluation of SARS-CoV-2 infected mouse lung, STable 3: Histological evaluation of SARS-CoV-2 infected hamster lung and STable 4: Histological evaluation of H1N1 infected mouse lung.

## Author Contributions

Conceptualization, Nicole Froelich, Girish Kirimanjeswara, Troy Sutton and Margherita Cantorna; Formal analysis, Juhi Arora, Jingpeng Wang, Erin Luley and Kathleen Kelly; Funding acquisition, Margherita Cantorna; Investigation, Juhi Arora, Devanshit Patel, McKayla Nicol, Cassamdra Fields, Katherine Restori, Jingpeng Wang, Nicole Froelich, Bhuvana Katkere, Josey Terwilliger, Veronika Weaver, Erin Luley, Kathleen Kelly and Margherita Cantorna; Methodology, Juhi Arora, Devanshit Patel, McKayla Nicol, Cassamdra Fields, Katherine Restori, Jingpeng Wang, Nicole Froelich, Bhuvana Katkere, Josey Terwilliger, Veronika Weaver, Erin Luley and Kathleen Kelly; Project administration, Juhi Arora, Devanshit Patel, McKayla Nicol, Cassamdra Fields, Bhuvana Katkere, Veronika Weaver, Girish Kirimanjeswara, Troy Sutton and Margherita Cantorna; Resources, Girish Kirimanjeswara, Troy Sutton and Margherita Cantorna; Supervision, Katherine Restori, Bhuvana Katkere, Girish Kirimanjeswara and Troy Sutton; Visualization, Juhi Arora, McKayla Nicol, Jingpeng Wang and Nicole Froelich; Writing – original draft, Juhi Arora and Margherita Cantorna; Writing – review & editing, Juhi Arora, Devanshit Patel, McKayla Nicol, Cassamdra Fields, Katherine Restori, Josey Terwilliger, Veronika Weaver, Kathleen Kelly, Girish Kirimanjeswara, Troy Sutton and Margherita Cantorna.

All authors will be informed about each step of manuscript processing including submission, revision, revision reminder, etc. via emails from our system or assigned Assistant Editor.

## Institutional Review Board Statement

All studies with SARS-CoV-2 were conducted in a biosafety level 3 enhanced (BSL3+) laboratory. This facility is approved for BSL3+ respiratory pathogen studies by the US Department of Agriculture and Centers for Disease Control. Studies with pandemic H1N1 influenza were conducted under biosafety level 2 enhanced conditions. All animal studies were conducted in compliance with the Animal Care and Use Committee under protocol numbers: 202001693, 202001516, 202001440 and 202001638.

## Informed Consent Statement

Not applicable.

## Data Availability Statement

Not applicable.

