## Supplementary data for "Vitamin D and the ability to produce 1,25(OH)_2_D are critical for protection from viral infection of the lungs"

**Supplementary Table 1. Primer sequences for qPCR.**

| **Target gene** | **Forward Primer (5’-3’)** | **Reverse Primer (5’-3’)** |
| --- | --- | --- |
| *Vdr* | 5’-CTG CAC CTC CTC ATC TGT GA-3’ | 5’-CCC CTT CAA TGG AGA TTG C-3’ |
| *Cyp24A1* | 5’-ACC CCC AAG GTC CGT GAC ATC-3 | 5’-CCA GTT GGG TCC AGG TAA GG-3’ |
| *Cyp27B1* | 5′-CCG CGG GCT ATG CTG GAA C-3′ | 5′-CTC TGG GCA AAG GCA AAC ATC TGA-3′ |
| *Ifnα* | 5'-GGA CTT TGG ATT CCC GCA GGA GAA G-3' | 5'- GCT GCA TCA GAC AGC CTT GCA GGT C-3' |
| *Ifnβ* | 5’-AGG GCG GAC TTC AAG ATC-3’ | 5’-CTC ATT CCA CCC AGT GCT-3’ |
| *Ifnγ* | 5’-TGC ATC TTG GCT TGG CAG CTC TTC-3’ | 5’-GGG TTG ACC TCA AAC TTG GCA-3’ |
| H1N1 M gene | 5’-AGA TGA GTC TTC TAA CCG AGG TCG-3’ | 5’-TCG AGA TCG GTG TTC TTT CC-3’ |

**Supplementary Table 2.** **Histological evaluation of SARS-CoV-2 infected mouse lung.^1^**

| **Parameter** | **Score = 0** | **Score = 1** | **Score = 2** | **Score = 3** |
| --- | --- | --- | --- | --- |
| **Perivascular Infiltrates**  **(0-3)** | None | Up to 10% vessels affected | Up to 25% vessels affected | ≥ 25% vessels affected |
| **Lymphocyte dominant**  **(0-1)** | No | Yes |  |  |
| **Endothelial reactivity**  **(0-2)** | None | Any/multi-focal | Generalized |  |
| **Other vascular parameters**  **(0-1)** | Absent | Present |  |  |
| **Type II pneumocyte hypertrophy**  **(0-1)** | Absent | Present |  |  |
| **Pneumonia extent**  **(0-3)** | None | Minimal, focal | Minimal, multi-focal | Mild, multi-focal |
| **Alveolar remodeling**  **(0-1)** | None | Any |  |  |
| **Interstitial pneumonia**  **(0-3)** | None | Up to 10% affected | 10- 25% affected | ≥ 25% affected |
| **Intra-alveolar inflammation (0-3)** | None | Up to 10% affected | Up to 25% affected | ≥ 25% affected |

^1^Histopathology criteria for scoring lung sections of SARS-CoV-2 infected mice.

**Supplementary Table 3.** **Histological evaluation of SARS-CoV-2 infected hamster.^1^**

| **Parameter** | **Score = 0** | **Score = 1** | **Score = 2** | **Score = 3** | **Score = 4** |
| --- | --- | --- | --- | --- | --- |
| **Lesions**  **(0-4)** | None | Up to 25% affected | 25-50% affected | 50-75% affected | > 75% affected |
| **Alveoli**  **(0-3)** | None | Mild edema and/or infiltrate | Moderate to severe edema or infiltrate | Diffuse alveolar damage | N/A |
| **Bronchioles**  **(0-3)** | None | Mild peribronchiolar cuffing and/or infiltrate, or epithelial degeneration | Moderate peribronchiolar cuffing and/or infiltrate with epithelial degeneration | Severe peribronchiolar cuffing and/or infiltrate with epithelial necrosis | N/A |
| **Blood vessels (0-3)** | None | Perivascular edema, mild perivascular cuffing | Perivascular cuffing in >30% of vessels, minimal vasculitis | Vasculitis with thrombosis | N/A |
| **Hemorrhage (0-2)** | None | Mild | Severe | N/A | N/A |
| **Type II pneumocyte hyperplasia (0-2)** | None | Early change, mild | Marked change, widespread | N/A | N/A |

^1^Histopathology criteria for scoring lung sections of SARS-CoV-2 infected hamsters.

**Supplementary Table 4.** **Histological evaluation of H1N1 infected mice.^1^**

| **Parameter** | **Score = 0** | **Score = 1** | **Score = 2** | **Score = 3** |
| --- | --- | --- | --- | --- |
| **Total lymphocyte infiltration**  **(0-3)** | None | Some inflammation | < 3 foci of inflammation | > 3 foci of inflammation |
| **Perivascular infiltration**  **(0-3)** | None | Some lymphocytes around bronchiole or blood vessel | Lymphoid aggregates around bronchioles or blood vessel | Multiple lymphoid aggregates |
| **Alveolar Hemorrhage**  **(0-3)** | None | Few hemorrhagic lesions | Multiple lesions | Hemorrhage with alveolar damage |

^1^Histopathology criteria for scoring lung sections of H1N1 influenza infected mice.
